# EasyMultiProfiler: An Efficient Multi-Omics Data Integration and Analysis Workflow for Microbiome Research

**DOI:** 10.1101/2025.04.17.649266

**Authors:** Bingdong Liu, Yaxi Liu, Shuangbin Xu, Qiusheng Wu, Dan Wu, Li Zhan, Yufan Liao, Yongzhan Mai, Minghao Zheng, Shenghe Wang, Yixin Chen, Zhipeng Huang, Xiao Luo, Zijing Xie, Xiaohan Pan, Guangchuang Yu, Liwei Xie

**Affiliations:** Zhujiang Hospital, Southern Medical University, Guangzhou, China; State Key Laboratory of Applied Microbiology Southern China, Guangdong Provincial Key Laboratory of Microbial Culture Collection and Application Guangdong Open Laboratory of Applied Microbiology, Institute of Microbiology, Guangdong Academy of Sciences, Guangzhou, China; Department of Bioinformatics, School of Basic Medical Sciences, Southern Medical University, Guangzhou, China; School of Life and Health Sciences, Fuyao University of Science and Technology, Fuzhou, China; Department of Psychiatry, The First Affiliated Hospital of Jinan University, Guangzhou, China; State Key Laboratory of Environmental Geochemistry, Institute of Geochemistry, Chinese Academy of Sciences, Guiyang, China; Pearl River Fisheries Research Institute, Chinese Academy of Fishery Sciences, Guangzhou, China; Department of Applied Biology and Chemical Technology, The Hong Kong Polytechnic University, Hung Hom, Kowloon, Hong Kong; Fuyang Institute of Zhejiang Chinese Medical University, Hangzhou, China

**Keywords:** Multi-Omics, Microbiome, Transcriptome, Metabolome, Integrative Analysis

## Abstract

Graphical Abstract

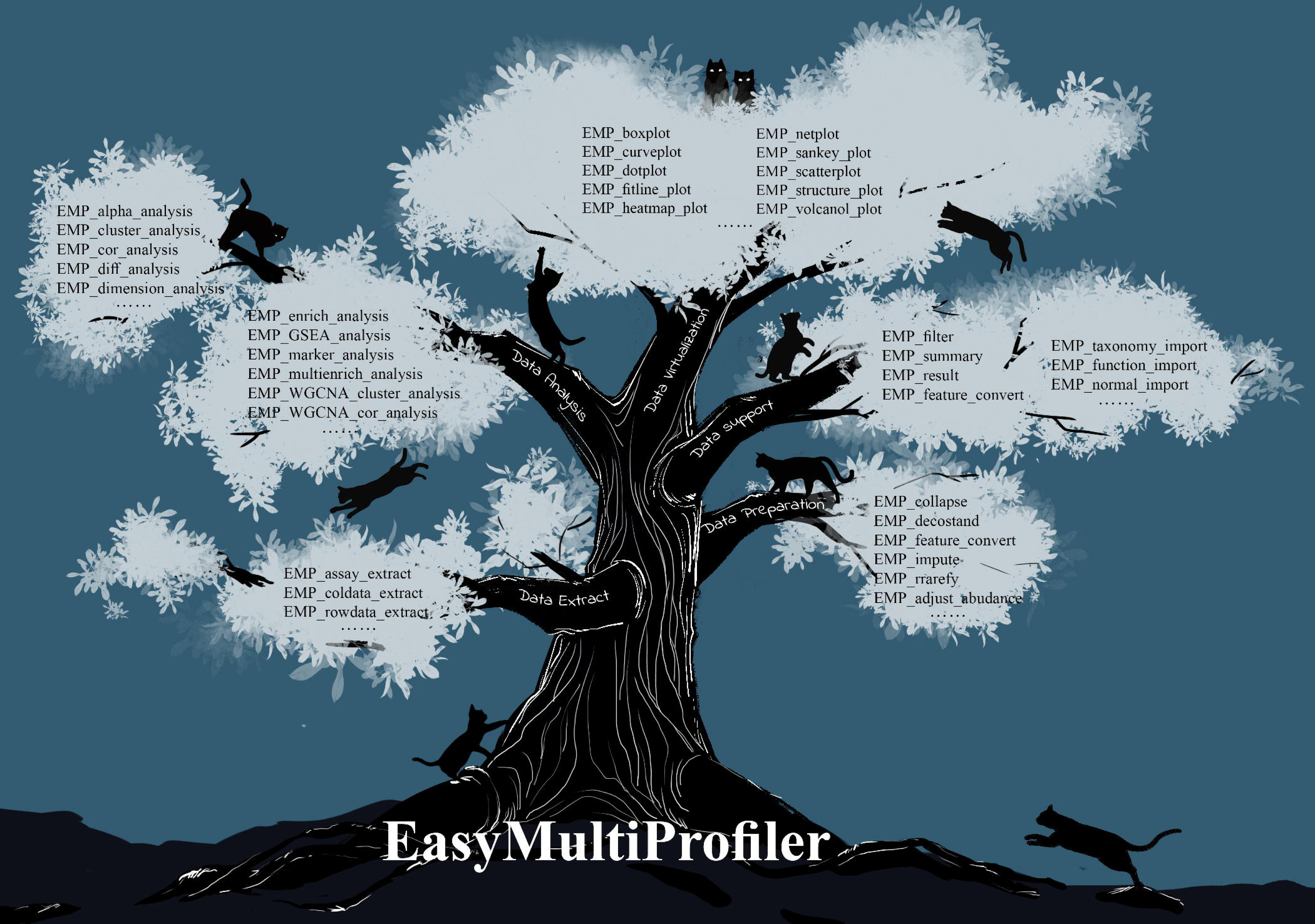

**Highlights:**

- EasyMultiProfiler provides unified integration of microbial multi-omics data
- EasyMultiProfiler enables natural language-style analytical workflows
- EasyMultiProfiler solves key challenges in standardization and reproducibility
- EasyMultiProfiler extracts meaningful biological insights efficiently

Host-microbiome interactions are crucial in maintaining physiological homeostasis and influencing disease progression. While traditional microbiome research, focused on microbial diversity and abundance, provided essential frameworks, the integration of multi-omics approaches established a far more comprehensive, systems-level understanding of microbiome functionality. However, significant methodological challenges in multi-omics data integration remain, including inconsistent sample coverage, heterogeneous data formats, and complex downstream analytical workflows. These challenges impact the reproducibility and reliability of results and underscore the critical requirement for developing standardized, systematic workflows. Thus, we developed the EasyMultiProfiler (EMP) workflow, a streamlined and efficient analytical framework for multi-omics data analysis. EMP provides a comprehensive infrastructure based on the SummarizedExperiment and MultiAssayExperiment classes, creating a unified framework for storing and analyzing omics data. The framework’s architecture comprises five interconnected functional modules: data extraction, preparation, support, analysis, and visualization. These modules are smoothly integrated into natural language-style analytical workflows, offering users an efficient and standardized solution for multi-omics analysis. EMP addresses critical challenges in multi-omics data analysis, including data integration, workflow standardization, and result reproducibility. The platform’s modular architecture and intuitive interface provide researchers and clinicians with a robust, flexible workflow to systematically extract biologically relevant insights from complex multi-omics datasets.

## Background

Recent development and progress in microbiome research have revealed extensive associations between microbial communities and host health(Lin et al. 2025). Accumulating evidence demonstrates that microbes not only participate in host physiological and metabolic processes but also play critical roles in the diseases’ development and progression(Agirman and Hsiao 2021; Liu et al. 2019). Thus, this understanding has placed microbiome research in a prominent scientific field, with increasing focus on elucidating molecular mechanisms underlying host-microbe interactions. Multi-omics approaches have emerged as the preferred methodological approaches for investigating this complex biological systems(Bai et al. 2025). However, multi-omics studies present significant challenges in data integration, methodological inconsistencies, and computational reproducibility, impeding the advancement of microbial multi-omics research (Figure 1).

**Figure 1.**
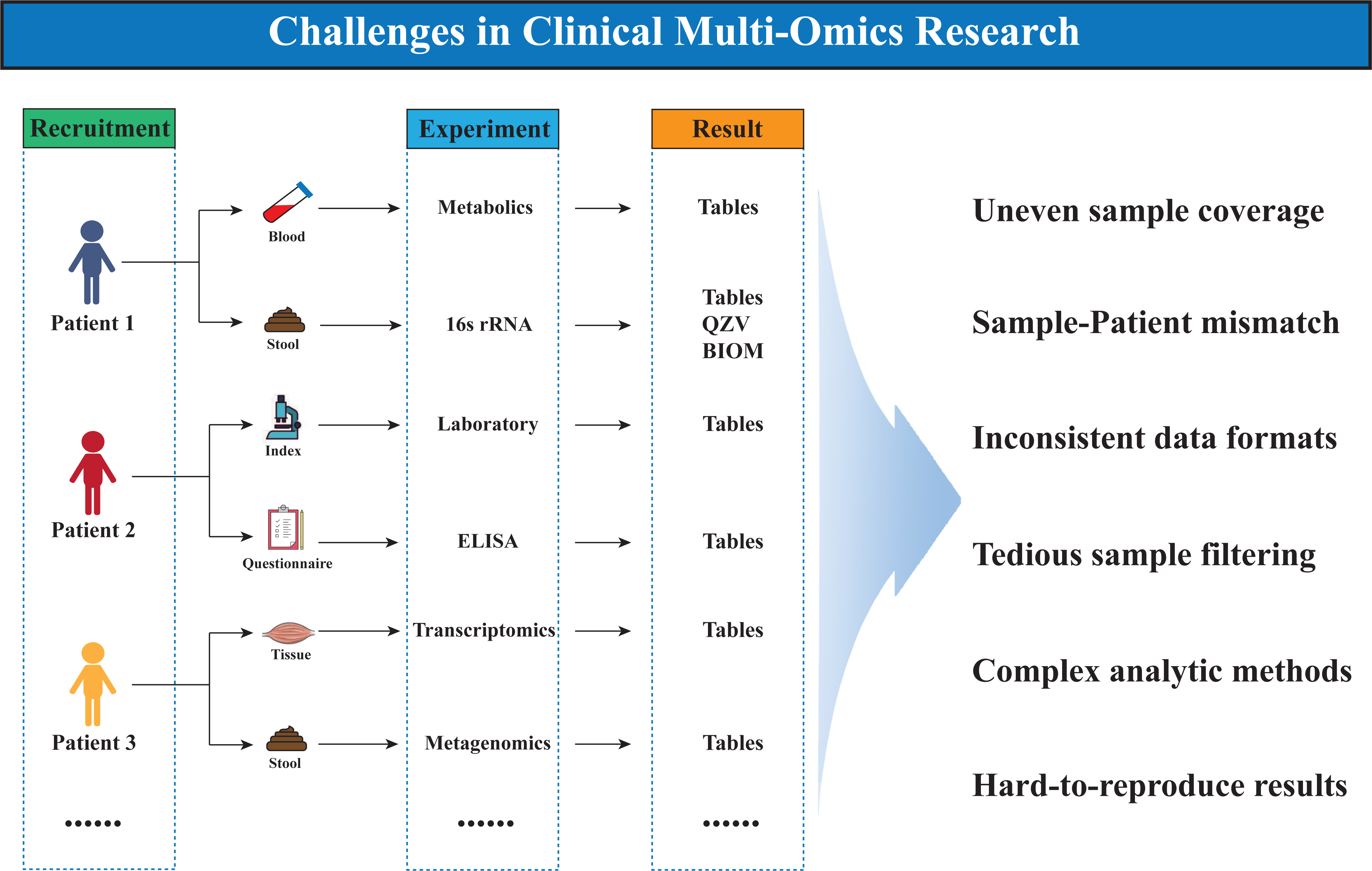
Clinical multi-omics research faces significant challenges, including reproducibility issues, data integration complexities, and analytical bottlenecks, which hinder a comprehensive understanding of host-microbiome interactions.

The primary challenge in front of us is the technical barriers regarding the data integration. Multi-omics datasets, e.g. microbiome, transcriptomics, and metabolomics frequently exhibit heterogeneous data structure and incomplete sample coverage. For example, microbial data are commonly stored in various formats, such as The Biological Observation Matrix (BIOM) from Mothur software, QIIME Zipped Visualization (qzv) files from QIIME2, or Metaphlan’s hierarchical annotation tables, while transcriptomic and metabolomic data are typically represented as annotated text files(Blanco-Míguez et al. 2023; Bolyen et al. 2019). This heterogeneity, combined with idiosyncratic metadata documentation practices, necessitates substantial computational effort for data processing and integration(Knight et al. 2018). Consequently, researchers must efficiently identify well-matched samples across all omics datasets during the analysis process, presenting a significant challenge for those without advanced data processing experience. Secondly, methodological selection and implementation present significant complexities and challenges as diverse analytical and bioinformatic toolkits exist for specific tasks. For example, microbial studies require rigorous preprocessing protocols, including normalization and filtering of low-abundance taxa, followed by the subsequent application of statistical and machine learning approaches (e.g., Random Forest, LASSO, XGBoost) for feature prioritization(Breiman 2001; Liu et al. 2022). In transcriptomic analysis, key genes are often identified using differential expression tools (e.g. DESeq2, limma, and edgeR) or advanced network-based methodologies such as WGCNA for identifying hub genes, while pathway-level interpretation is typically performed using enrichment analysis frameworks, such as Over-Representation Analysis (ORA) and Gene Set Enrichment Analysis (GSEA) (Langfelder and Horvath 2008; Love et al. 2014; Ritchie et al. 2015; Xu et al. 2024). However, the combination of different analytical pipelines with data preprocessing, normalization, and dimensionality reduction approaches even generates conflicting results(Li 2022; Nearing et al. 2022). The reproducibility of research findings remains a critical concern in multi-omics studies(He et al. 2018). Thirdly, beyond the inherent variability of microbial communities, the lack of transparent analytical workflows and standardized computational frameworks also hinders result validation(Schloss 2018). In most multi-omics analyses, the complexity of analytical pipelines—particularly the integration of diverse tools—along with inconsistencies in code functions, formatting, and the lack of standardized coding practices among bioinformaticians severely undermines the readability of code. Furthermore, the intricate nature of multi-omics analyses, combined with inconsistent implementation practices, creates substantial obstacles to research reproducibility and the effective transfer of knowledge within the biomedical field.

To address these challenges, we developed the EasyMultiProfiler (EMP) workflow, a streamlined and efficient analytical framework for multi-omics data analysis, particularly for integrative analysis of microbiome, transcriptomics, and metabolomics datasets. EMP provides a comprehensive solution for data import, processing, and visualization, while emphasizing computational reproducibility and methodological standardization, enabling researchers and clinicians to efficiently identify biological targets from complex multi-omics datasets.

## Method

### Data Container Architecture

The EMP framework implements a robust data container architecture utilizing the MultiAssayExperiment class for multi-omics data integration, complemented by the EMPT class for single-omics data management (Ramos et al. 2017). The EMPT class inherits from the SummarizedExperiment class, storing the abundance matrix, feature annotations, and phenotypic data while incorporating multiple parameter slots to streamline the workflow (Figure S1A)(Morgan et al. 2020). For integrative analysis, EMPT architecture enables dataset merging through an intuitive “+” operator (Figure S1B). The R package was constructed using S4 syntax to ensure data encapsulation, thereby preventing unintended modifications during user operations. Furthermore, the container architecture incorporates well-defined method interfaces, significantly enhancing compatibility and extensibility for subsequent development.

### Data Import Framework

To address heterogeneous formats of multi-omics data, EMP implements optimized import functions for multi-omics through a one-step conversion into the data container. For microbial data, the EMP_taxonomy_import function processes microbial taxonomic data from multiple platforms (e.g., qzv files, and biom formats generated by tools such as MetaPhlAn3, QIIME2, and Mothur), automatically parsing taxonomic hierarchical taxonomic structures. For functional annotations, the EMP_function_import processes KO/EC annotation data generated by PICRUst2 or HUMAnN2, with automatic annotation completion based on the KEGG database(Beghini et al. 2021). For transcriptomic and metabolomic data, the EMP_normal_import function was designed to efficiently extract annotation information and abundance matrices from a standardized annotation table. Following the import of individual omics datasets, users could easily integrate them into the MultiAssayExperiment container according to metadata and sample map information (Figure 2).

**Figure 2.**
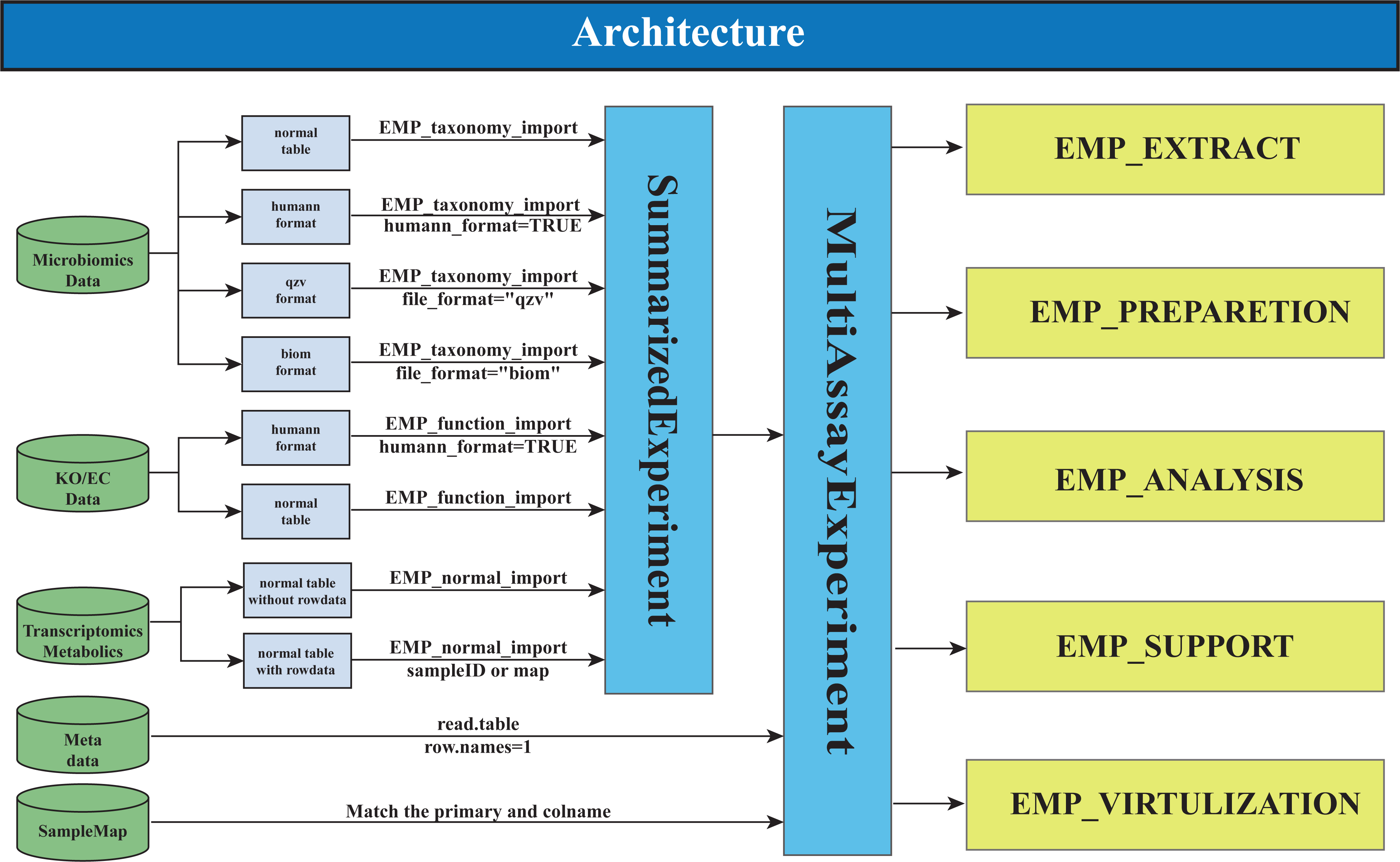
The architecture of The EasyMultiProfiler package.

### Modular Analytical Workflow

EMP workflow implements five core modules: Data Extraction, Data Preparation, Data Analysis, Data Support, and Data Visualization, offering comprehensive multi-omics analysis capabilities with a streamlined and efficient design (Figure 3A, B). The Data Extraction module enables either selective omics data extraction for downstream analysis or meta-information-based multi-omics datasets filtering, exporting them in the MultiAssayExperiment format to evaluate potential research projects. The Data Preparation incorporates normalization, batch effect removal, data collapsing, and feature ID conversion to help users prepare their data for formal analysis. The Data Analysis module integrates established methodologies for differential analysis, dimensionality reduction, machine learning, enrichment analysis, network, etc. The Data Visualization module provides corresponding visualization capabilities for each analytical component, eliminating the need for manual data interfacing and ensuring a streamlined handling experience. Additionally, the Data Support module enables data filtering and analysis history tracking (Figure 3C, Figure S2A). Following tidyverse design principles, EMP analytical workflow utilizes pipe operators (|> or %>%) for seamless module integration, enhancing code readability and analytical reproducibility and providing users with an efficient and intuitive experience for multi-omics data analysis.

**Figure 3.**
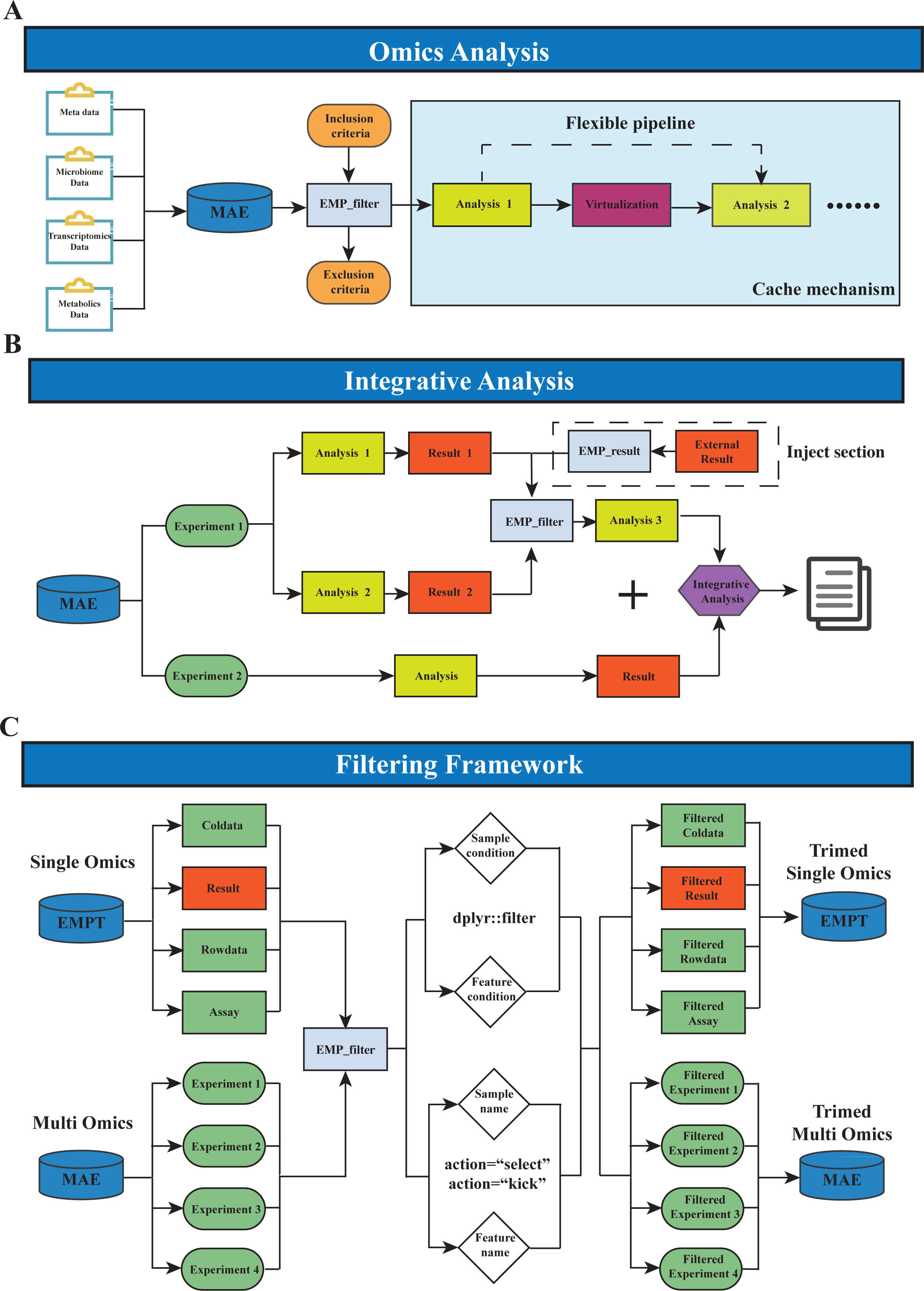
The mechanism of the EasyMultiProfiler (EMP) workflow for omics data analysis. (A, B) The workflow of EMP for processing clinical multi-omics data and performing integrative analysis. (C) The mechanism of EMP for data and result filtering.

### User-friendly design

EMP is designed for efficient and user-friendly multi-omics analysis. Its module-level caching minimizes redundant computations (Figure S2B). A simple “+” operator simplifies dataset merging for multi-omics integration, making tasks like correlation analysis and Weighted Gene Co-Expression Network Analysis (WGCNA) more straightforward. The EMP_filter function allows flexible data and result filtering at any point, with automatic suitability checks and the creation of filtered data containers (Figure S1C). EMP_result facilitates integration with other R packages by importing their results into the EMP workflow. EMP_save_var allows users to save current results as environment variables without interrupting the analysis. Finally, more utility functions like str_detect_multi and top_detect improve analytical efficiency by simplifying feature manipulation.

### Comprehensive tutorials

The complete source code is publicly available on GitHub (https://github.com/liubingdong/EasyMultiProfiler), accompanied by a comprehensive and step-by-step online documentation (http://easymultiprofiler.xielab.net/) providing detailed tutorials for the complete analytical workflow.

## Result

To demonstrate the EMP’s analytical capacities, we applied the workflow to two large-scale omics datasets: (1) diet-microbiome interaction in patients with metabolic syndrome and (2) gut microbiome-metabolome associations in patients with colorectal cancer.

### Example 1: Diet-Microbiome Interaction in Patients with Metabolic Syndrome Based on Large-Scale Population

Utilizing the Guangdong Gut Microbiome Project dataset (GGMP), we analyzed associations among gut microbiota, dietary patterns, and clinical phenotypes in patients with metabolic syndrome. Following microbial data in BIOM format import into EMP workflow by generating a MultiAssayExperiment object (Figure 4A), dietary pattern analysis of 7,009 samples revealed 6,703 samples (95.6%) with consistent dietary behaviors (Figure 4B, Tables S1-S2). After excluding subjects with confounding conditions (antibiotic use, colitis, or diarrhea/constipation), 5388 individuals remained including 1,080 with metabolic syndrome.

**Figure 4.**
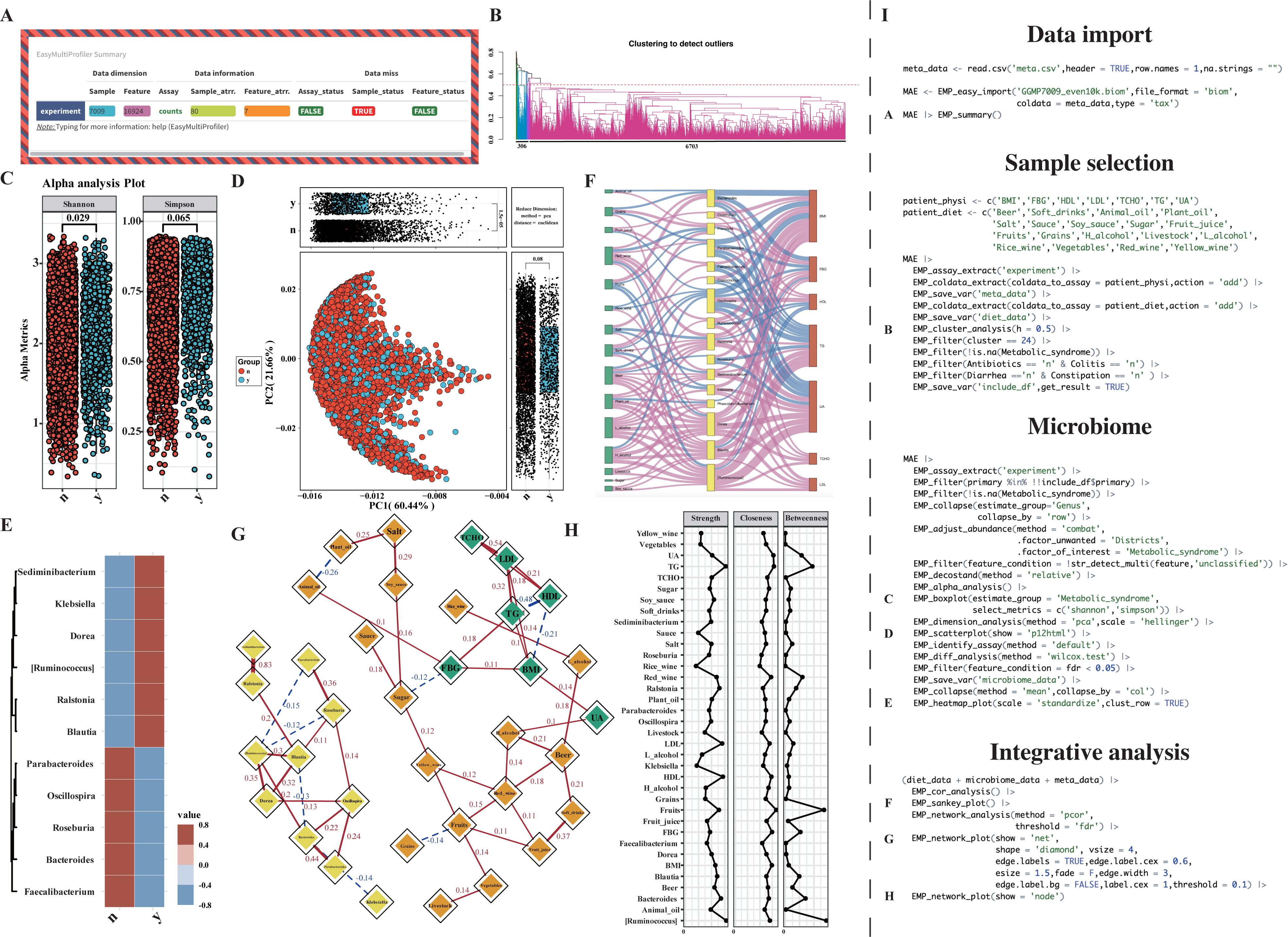
Investigating the Relationship Between Diet and Gut Microbiome in Metabolic Syndrome Patients. (A) EMP package could import microbial data in BIOM format. (B) Clustering analysis of dietary data from 7,009 samples revealed that 6,703 samples (95.6%) exhibited similar dietary patterns. (C, D) Alpha and beta diversity analyses demonstrated significant differences between groups by Wilcoxon test, where “y” represents the metabolic syndrome group and “n” denotes the control group. (E) Heatmap displaying differential taxa identified by Wilcoxon test (FDR < 0.05). (F) Sankey plot based on Spearman correlation analysis illustrated interactions among diet, microbiome, and disease-related indices. (G, H) Partial correlation network and three algorithms for evaluating node importance. (I) The code for all the figures above.

Considering that regional factors are the most significant influence on microbial composition other than disease, we applied the combat algorithm to correct batch effects in the microbial abundance matrix and perform analyses at the genus level. After filtering out taxa with missing exact annotations, 752 genera were retained for downstream analysis. Alpha diversity analysis revealed that the diversity in the metabolic syndrome group was different from that in the non-metabolic syndrome group (Shannon *pvalue* = 0.029; Simpson *pvalue* = 0.065) (Figure 4C, Supplementary Tables S3). Principal component analysis (PCA) dimensionality reduction further confirmed significant differences in microbial community structure between the two groups (Figure 4D). According to the Wilcoxon test, we identified 11 genera with significant differences between the groups (FDR < 0.05) (Figure 4E, Supplementary Tables S4).

To explore the complex relationships among diet, microbiome, and disease phenotypes, we first constructed a Sankey diagram based on Spearman correlation, visually illustrating the complex interactions among the three data (Figure 4F). Given the high collinearity among multi-omics data, we further constructed a partial correlation network (PCOR) and revealed that features from the three omics tightly constructed a complex interaction network (Figure 4G, Supplementary Tables S5). According to three algorithms of evaluating the importance of nodes, we found that, at the dietary level, fruit intake had the most significant impact on the network; at the microbial level, the genera *Ruminococcus* and *Bacteroides* occupied central positions; and at the disease phenotype level, triglyceride (TG) levels exhibited the strongest network influence (Figure 4H, Supplementary Tables S6).

### Example 2: Gut Microbiome-Metabolome Associations in Patients with Colorectal Cancer Based on Multi-Study Data

We collected gut microbiome and metabolome data from 1,483 individuals across five studies, including 480 healthy subjects (HS group), 48 patients with multiple polyp adenomas (MP group), 305 patients with irritable bowel syndrome (IBS group), 478 patients with inflammatory bowel disease (IBD group), 150 patients with colorectal cancer (CRC group), and 30 normal individuals with a history of colorectal surgery (NHS group). After batch effect correction, we observed significant differences in microbial diversity at the genus level between the control group and all other groups except the MP and IBD groups (Figure 5A, Supplementary Tables S7). Dimensionality reduction analysis further revealed that, along with the situation of intestinal disease severity, the microbial community structures of most groups differed significantly from that of the control group (Figure 5B). Interestingly, the microbial structure in the NHS group is similar to that of the control group, suggesting that the gut microbiota of postoperative colorectal cancer patients may have progressively recovered to a healthy state. To further discover the taxa maker, we employed the Boruta algorithm to identify 27 genera with potential biological significance among groups (Supplementary Tables S8). Notably, *Mediterraneibacter* was significantly abundant in the CRC group compared to other groups (Figure 5C).

**Figure 5.**
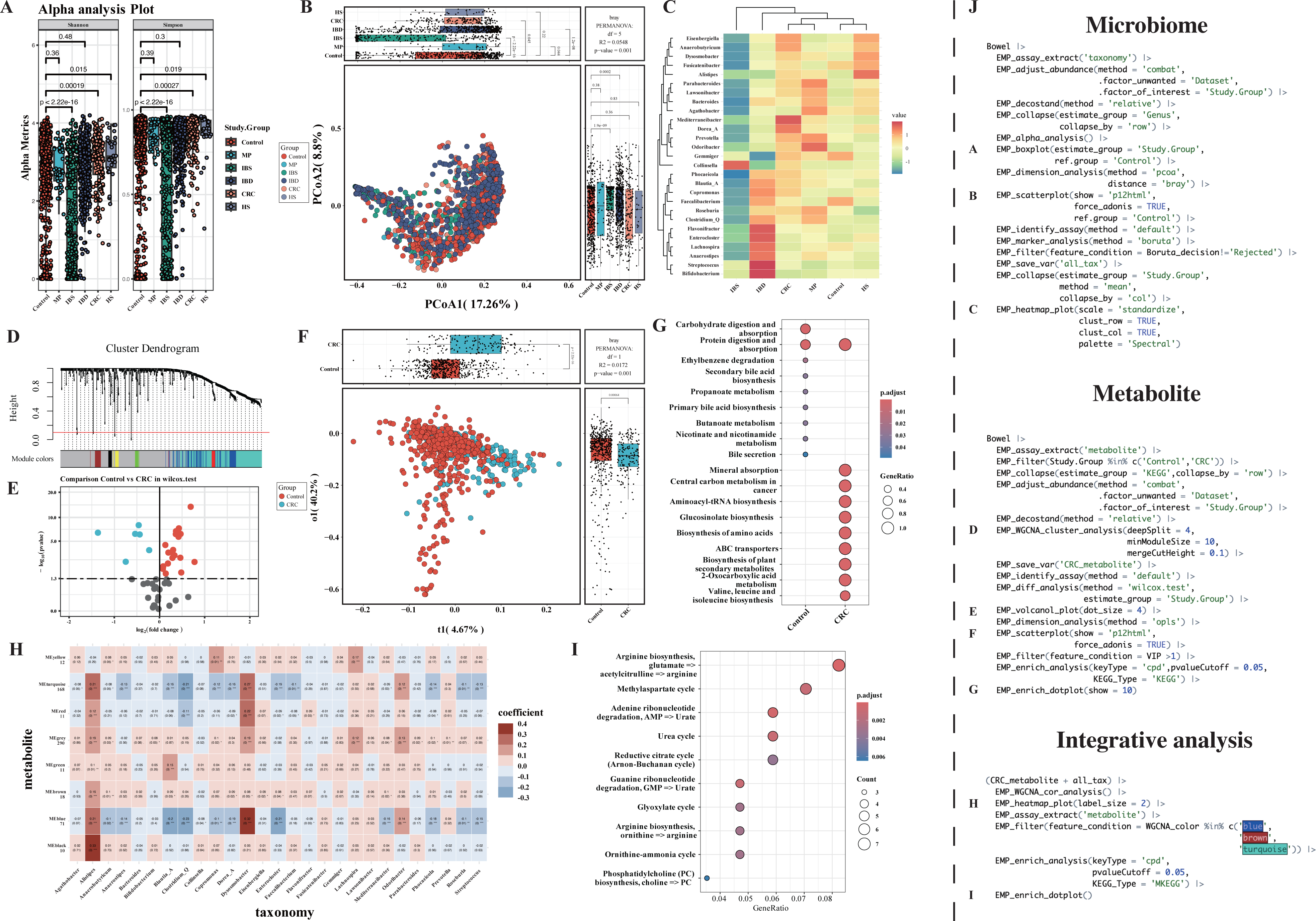
Integrative Analysis of Microbiome and Metabolome in Colorectal Cancer Based on Multi-study Data. (A, B) Alpha and beta diversity analyses demonstrated significant differences between the control and CRC group by Wilcox and Permutational multivariate analysis of variance (PERMANOVA). (C) Boruta algorism identified the marker taxa among groups. (D) The WGCNA cluster analysis identified 8 modules from metabolite data. (E, F) OPLS and differential analysis showed metabolite structure alteration in the CRC group. (G) The differential metabolites (VIP >1) contributed to many cancer-related pathways in KEGG enrichment. (H, I) The WGCNA analysis showed the differential metabolites associated with *Mediterraneibacter* contributed to arginine-related metabolic pathways. (J) The code for all the figures above.

At the metabolome level, after batch effect correction, orthogonal projections to latent structures (OPLS) analysis demonstrated significant differences in metabolite profiles between the CRC group and the control group (Figure 5E, F, Supplementary Tables S9). By screening metabolites with VIP > 1 threshold and performing KEGG pathway enrichment analysis, we found that the differential metabolites in the CRC group were significantly enriched in pathways that were associated with the cancer, such as “central carbon metabolism in cancer” and “aminoacyl-tRNA biosynthesis”, suggesting that gut microbiota may participate in the pathophysiology of CRC by modulating these metabolic pathways (Figure 5G, Supplementary Tables S10).

To explore the crosstalk between microbial biomarkers and metabolites, we employed the WGCNA strategy to estimate the association between metabolites and *Mediterraneibacter*. The metabolites clustered into eight modules, and three modules, including 257 metabolites, showed significant correlations with *Mediterraneibacter* (Figure 5D, H). KEGG enrichment showed these metabolites primarily involved in the arginine metabolism-related pathways (Figure 5I, Supplementary Tables S11).

## Discussion

### Current Challenges in Multi-omics Research

Microbial single-omics data only consists of three components: abundance table (microbial abundance matrices), feature annotations (taxonomic annotation tables), and sample metadata (clinical phenotype data)(Liu et al. 2020). However, microbial multi-omics research faces significant challenges in managing complex and multidimensional data structures across multiple layers, such as microbial abundance, functional genes, metabolomes, transcriptomes and phenotypical data. This cross-omics data integration not only requires the coordination of complex data files (e.g., sample matching and data format standardization) but also addresses the dynamic interrelationships among multi-omics datasets. However, traditional data management approaches use manual operations on tabular files (e.g., Excel or CSV) for data matching and format conversion, which is inefficient and inadequate for large-scale studies, particularly with coordinating multi-omics data integration, sample matching and format standardization. The shift from hypothesis-driven to data-driven research paradigms further emphasizes these limitations, highlighting the need for more robust data management solutions.

### EMP’s Innovative Framework

EMP workflow addresses these challenges through three key innovations. First, at the data structure level, EMP leverages SummarizedExperiment and MultiAssayExperiment frameworks to construct standardized data containers. This design not only inherits the universal data architecture widely adopted in bioinformatics but also achieves self-consistent storage of multi-omics data through object-oriented programming. Each omics dataset independently encapsulates abundance matrices, feature annotations, and sample metadata, while enabling cross-omics associations via unified interfaces. Second, at the analytical workflow level, EMP’s modular design philosophy deconstructs complex analyses into functional units (data extraction-preprocessing-analysis-visualization), significantly reducing operational complexity. Third, at the level of methodological compatibility, the framework maintains methodological compatibility through the EMP_result interface, ensuring workflow stability while enabling expansion.

### Enhanced User Experience

Unlike traditional muti-omics tools (e.g. mixOmics or OmicsNet) that primarily focus on specific analytical steps but seldomly offer help to manage the whole workflow, which requires researchers to master complex skills (Table 1)(Chen et al. 2019; Ding et al. 2021; Langfelder and Horvath 2008; Tang et al.; Tebaldi et al. 2014; Welham et al. 2023; Zhou et al. 2022), EMP made significant optimization and improvement on user experience through three key strategies. First, EMP implements the tidyverse-like pipe operators for intuitive workflow construction (e.g., dataset |> normalize() |> differential_analysis() |> visualize()). Second, EMP redefines the “+” operator to combine omics data in the integrative analysis (e.g. microbiome + metabolome → network analysis). This design works more closely with intuition, enabling researchers to focus on addressing scientific questions rather than expending effort on technical intricacies. Third, the EMP workflow incorporates caching technology, allowing users to interactively refine their analysis strategies at each step based on intermediate results while minimizing redundant computational time and resource consumption (Figure S2A).

**Table 1.**
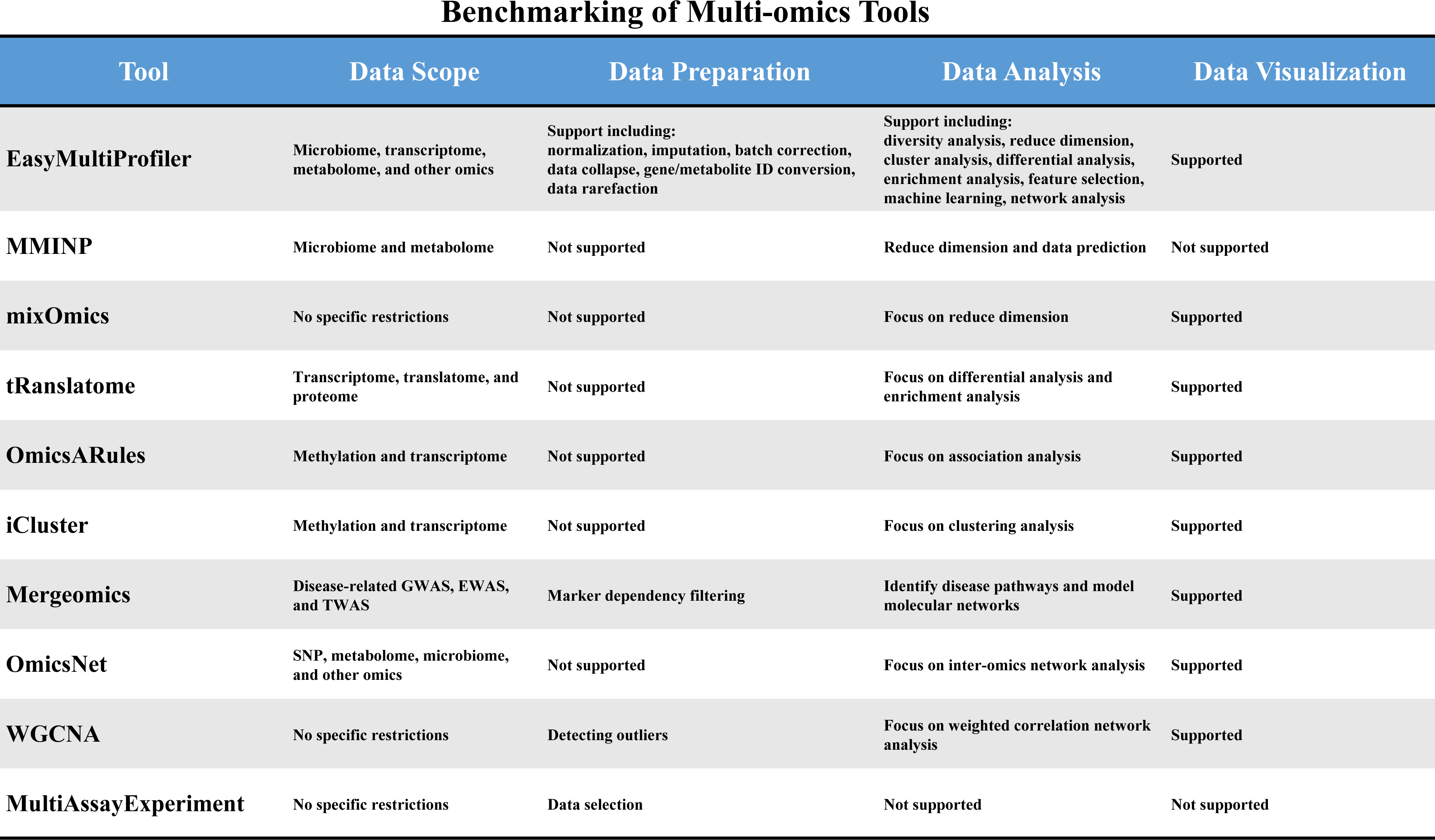
EasyMultiProfiler demonstrated comprehensive capabilities compared to existing multi-omics tools.

### Validation Through Real-world Applications

In the open-source study testing, the workflow’s reliability was thoroughly validated through two comprehensive case studies. In metabolic syndrome dataset () from the Guangdong Gut Microbiome Project (Example 1: n=7,004), EMP successfully analyzed the relationships among diet, microbiome, and disease phenotypes in patients with metabolic syndrome while controlling for dietary and regional factors and identified previously reported 11 genera with strong disease-association with metabolic syndrome(Chanda et al. 2024; Kuang et al. 2023; Munukka et al. 2017; Takeuchi et al. 2023). Notably, the genera *Ruminococcus* and *Bacteroides* played central roles in the partial correlation network across the three omics layers. Previous studies have shown that *Ruminococcus gnavus* is associated with various phenotypes such as obesity, allergies, and infections, suggesting its potential as a biomarker for metabolic syndrome(Hong et al. 2024). Meanwhile, *Bacteroides* has been found to enhance weight loss in subjects on low-carbohydrate diets, indicating its potential as a probiotics(Zhang et al. 2021). In the muti-study across five datasets (Example 2: n=1,483), the significant alterations in gut microbiota and metabolites in colorectal cancer (CRC) patients aligned with established literature on CRC pathophysiology, with a notable increase in the abundance of the genus Mediterraneibacter in the CRC group. Mediterraneibacter, a newly established genus, includes reclassified species previously belonging to Ruminococcus, such as Ruminococcus gnavus and Ruminococcus torques, which have been reported as biomarkers for colorectal cancer (CRC), and were associated with shortened colon length and exacerbated inflammatory responses(Huang et al. 2024; Togo et al. 2018; Wu et al. 2021). Further integrative omics analysis revealed that 257 differentially abundant metabolites closely associated with Mediterraneibacter were primarily enriched in arginine-related metabolic pathways. Previous studies have demonstrated that arginine promotes the proliferation of CRC tumor cells, and modulating the gut microbiota to reduce arginine synthesis has emerged as an effective strategy for CRC treatment(Alexandrou 2018; Jin et al. 2025; Liu et al. 2024). These two examples comprehensively demonstrate the efficiency, simplicity, practicality, and reliability of the EMP analytical workflow.

### Current Limitations

Despite the significant advantages demonstrated by EMP in multi-omics integration analysis, several current limitations must be acknowledged: (1) Limited support for emerging technologies (e.g. single-cell omics); (2) Suboptimal parallel computing efficiency for large-scale datasets; and (3) Absence of a graphical user interface (GUI). Future improvement priorities will focus on three key directions: (1) Expanding compatibility with other data types, such as single-cell and spatial transcriptomics data; (2) Enhancing computational efficiency for large-scale datasets; (3) Implementing a shiny-based visual programming interface. The establishment of an EMP open-source community will serve as a long-term strategy, continuously integrating cutting-edge analytical methods through collaborative efforts to sustain the tool’s vitality.

### Conclusion

EMP workflow represents a significant advance in microbial multi-omics research through its innovative architecture and optimized user experience. Its modular, standardized, and open framework not only significantly improves data analysis efficiency, but also establishes a traceable and reproducible research paradigm. As multi-omics technologies increasingly integrate into clinical diagnostics and precision medicine, the EMP workflow serves as a vital infrastructure, bridging fundamental research and translational applications while offering robust methodological support for unraveling the complex networks of microbe-host interactions.

## Availability of data and materials

The data for the two examples in this study were collected from the Gut Microbiome Project (https://github.com/SMUJYYXB/GGMP-Regional-variations) and the microbiome-metabolome dataset from the Borenstein Lab (https://github.com/borenstein-lab/microbiome-metabolome-curated-data). All code and raw data used in this study are publicly available (https://github.com/liubingdong/EasyMultiProfiler_paper_examples).

## Author contributions

LW.X. and GC.Y. designed the study and acquired grant support. BD.L. constructed the R package. BD.L. and YX.L. drafted the manuscript and tutorial. SB.X., QS.W. and L.Z. performed the code review. D.W, YF.L., MH. Z, SH. W and YX.C. collected the data. YZ.M, X.L., ZJ.X, and XH.P. performed unit test.

## Supporting Information

**Figure S1. Detailed information of EasyMultiProfiler.** (A) The structure of single-omics data consists of an abundance matrix, metadata, and feature information. (B) The EMPT class could be combined by operator “+”. (C) The filter function in the workflow could filter the results accordingly when samples and features change.

**Figure S2. Demonstration of the caching and history functions in the EMP workflow.** (A) The history function enables detailed tracing of results or visualizations at any step within a complex workflow. (B) The caching function is automatically activated, significantly reducing computational time for repeated steps.

