## Supplementary Figure 1 for "EasyMultiProfiler: An Efficient Multi-Omics Data Integration and Analysis Workflow for Microbiome Research"

A

| Assay | Experiment data | | | | | |
|---|---|---|---|---|---|---|
| | feature1 | feature2 | feature3 | feature4 | feature5 | …… |
| P1 | | | | | | |
| P2 | | | | | | |
| P3 | | | | | | |
| …… | | | | | | |

| Coldata | Sample related data | | | | | |
|---|---|---|---|---|---|---|
| | info_1 | info_2 | info_3 | info_4 | info_5 | …… |
| P1 | | | | | | |
| P2 | | | | | | |
| P3 | | | | | | |
| …… | | | | | | |

| Rowdata | Feature related data | | | | | |
|---|---|---|---|---|---|---|
| | info_1 | info_2 | info_3 | info_4 | info_5 | …… |
| feature1 | | | | | | |
| feature2 | | | | | | |
| feature3 | | | | | | |
| …… | | | | | | |

B

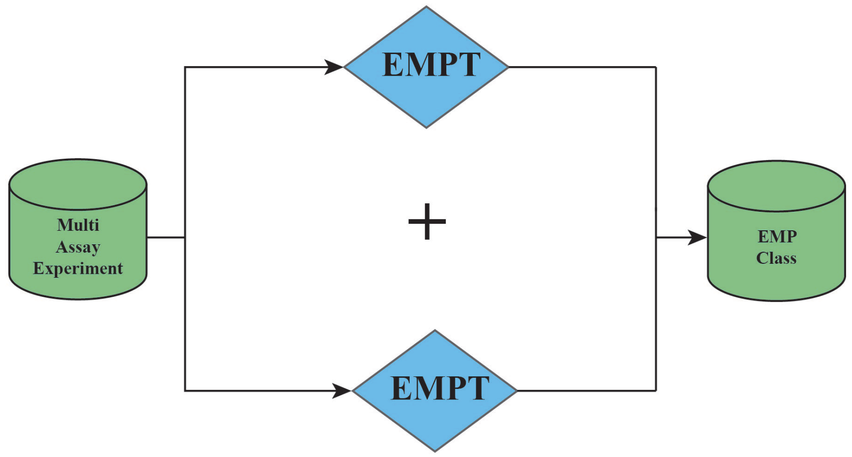


C

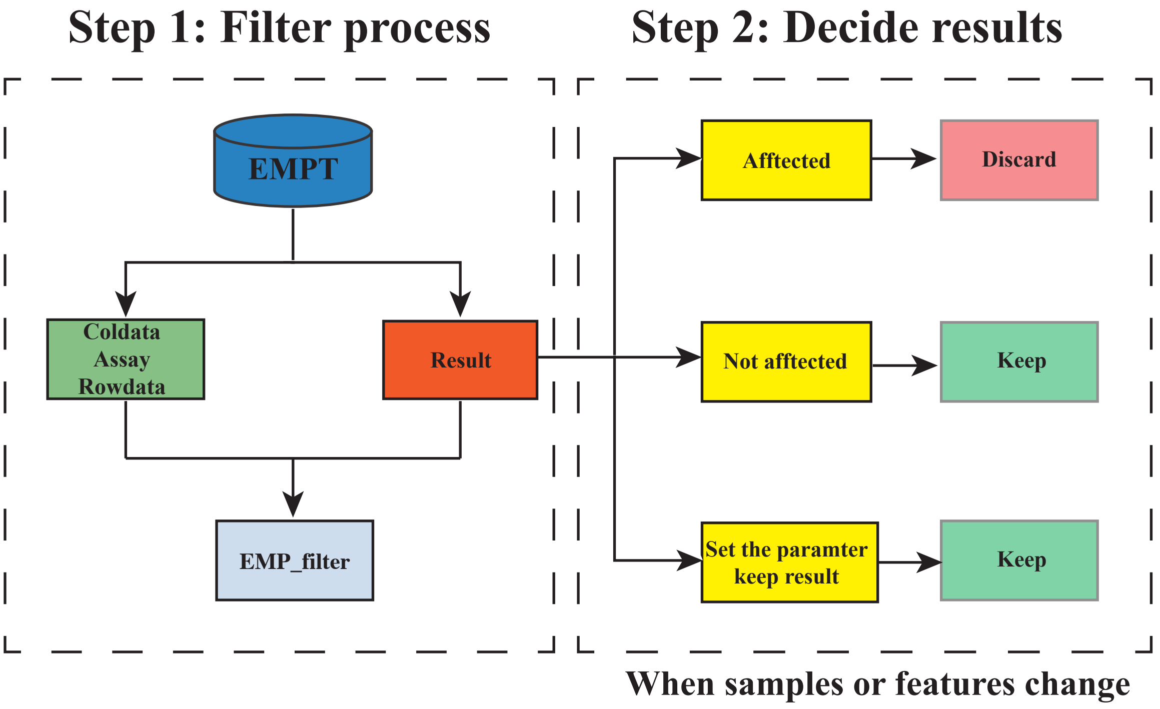
