## Supplementary Figure 2 for "EasyMultiProfiler: An Efficient Multi-Omics Data Integration and Analysis Workflow for Microbiome Research"

**A**

```r
library(magrittr)
library(EasyMultiProfiler)
data(MAE)
k1 <- MAE %>%
  EMP_assay_extract('taxonomy') %>%
  EMP_collapse(estimate_group = 'Genus',collapse_by = 'row') %>%
  EMP_diff_analysis(method='DESeq2', .formula = ~Group) %>%
  EMP_filter(feature_condition = pvalue<0.05)

k2 <- MAE %>%
  EMP_collapse(experiment = 'untarget_metabol',na_string=c('NA','null','','-'),
               estimate_group = 'MS2kegg',method = 'sum',
               collapse_by = 'row') %>%
  EMP_diff_analysis(method='DESeq2', .formula = ~Group) %>%
  EMP_filter(feature_condition = pvalue<0.05 & abs(fold_change) > 1.5)

#For two experinemnts
p1 <- (k1 + k2) %>% EMP_cor_analysis(method = 'spearman') %>%
  EMP_heatmap_plot()

p1 %>% EMP_history()
```

```
> p1 %>% EMP_history()
$total
[1] "as.EMP(object = data_list)"                    "EMP_cor_analysis(EMP = ., method = spearman)"
[3] "EMP_heatmap.EMP_cor_analysis(obj = obj)"

$taxonomy
[1] "EMP_assay_extract(obj = ., experiment = taxonomy)"
[2] "EMP_collapse(obj = ., estimate_group = Genus, collapse_by = row)"
[3] "EMP_diff_analysis(obj = ., .formula = ~Group, method = DESeq2)"

$untarget_metabol
[1] "EMP_collapse(obj = ., experiment = untarget_metabol, estimate_group = MS2kegg, method = sum, na_string = c(NA, null, , -), collapse_by = row)"
[2] "EMP_diff_analysis(obj = ., .formula = ~Group, method = DESeq2)"
```

**B**

```
> ## First run
> system.time({
+ MAE |>
+   EMP_assay_extract('geno_ec') |>
+   EMP_WGCNA_cluster_analysis(RsquaredCut = 0.85)
+ })
   user  system elapsed
  7.483   0.199   7.721
> ## Second run
> system.time({
+   MAE |>
+     EMP_assay_extract('geno_ec') |>
+     EMP_WGCNA_cluster_analysis(RsquaredCut = 0.85)
+ })
   user  system elapsed
  0.004   0.000   0.004
> ## Close the cached
> system.time({
+   MAE |>
+     EMP_assay_extract('geno_ec') |>
+     EMP_WGCNA_cluster_analysis(RsquaredCut = 0.85,use_cached = FALSE)
+ })
   user  system elapsed
  6.485   0.128   6.646
```
